# Deep-sea *in situ* and laboratory proteomics provide insights into the sulfur metabolism of a novel deep-sea bacterium, *Pseudodesulfovibrio serpens* sp. nov.

**DOI:** 10.1101/2023.10.05.561093

**Authors:** Chong Wang, Rikuan Zheng, Chaomin Sun

**Author notes:** Corresponding author: Chaomin Sun. E-mail address.

## Abstract

Sulfate-reducing bacteria (SRB) are ubiquitously distributed across various biospheres and play key roles in global sulfur cycles. However, few deep-sea SRB have been cultivated and studied *in situ*, limiting our understanding of the true metabolism of SRB in the deep biosphere. Here, we firstly clarified the high abundance of SRB in deep-sea cold seep sediments and successfully isolated a sulfate-reducing bacterium (strain zrk46). Our genomic, physiological and phylogenetic analyses indicate that strain zrk46 is a novel species, which we propose as *Pseudodesulfovibrio serpens*. Based on the combined results from growth assays and proteomic analyses, we found that supplementation with sulfate (SO_4_^2-^), thiosulfate (S_2_O_3_^2-^), or sulfite (SO_3_^2-^) promoted the growth of strain zrk46 by facilitating energy production through the dissimilatory sulfate reduction with the auxiliary functions of heterodisulfide reductases, ferredoxins, and nitrate reduction associated proteins, which were coupled with the oxidation of environmental organic matter in both laboratory and deep-sea *in situ* conditions. Moreover, metatranscriptomic results confirmed the dissimilatory sulfate reduction of deep-sea SRB in deep-sea environment, which might be coupled to the methane oxidation of anaerobic methanotrophic archaea (ANME-2) through direct interspecies electron transfer via cytochromes.

**IMPORTANCE:** The deep-sea cold seep sediments were ideal habitats for uncovering diverse metabolisms of SRB. Unfortunately, the paucity of SRB isolates has limited further insights into their physiological and metabolic features as well as ecological roles. In the present study, we demonstrated the high abundance of SRB in the deep-sea cold seep sediments and isolated a sulfate-reducing bacterium. Our results demonstrate that the existence of dissimilatory sulfate reduction of strain zrk46 in both laboratory and deep-sea *in situ* environments, accompanied by the auxiliary effect of heterodisulfide reductases, ferredoxins, and nitrate reduction associated proteins. Our findings also unravel that the sulfate reduction of deep-sea SRB in *in situ* environment might be coupled to the methane oxidation of ANME-2. Overall, these findings expand our understanding of deep-sea SRB, while highlighting their importance for deep-sea elemental cycles.

## INTRODUCTION

Sulfate-reducing bacteria are universally distributed in many engineered and natural environments where sulfate is present (1), which can obtain energy by oxidizing organic compounds or molecular hydrogen while reducing sulfate to hydrogen sulfide. Most SRB can also reduce other oxidized inorganic sulfur compounds, such as thiosulfate or elemental sulfur/polysulfide (2). It is known that SRB exist ubiquitously in deep oceans or terrestrial environments such as cold seeps, hydrothermal vents, oil reservoirs, aquifers, and groundwaters, where their geomicrobiological significance has often been emphasized (3, 4). The growth of these organisms is highly favored by an anaerobic (oxygen-free) environment. SRB were one group of sulfate reducing prokaryotes, and among the genera of SRB, the most thoroughly studied species are classified in the genus *Desulfovibrio* (5, 6), which is the most common sulfate-reducing organism that can obtain its energy predominantly from the anaerobic reduction of sulfates. Sulfate reduction includes two approaches: assimilatory sulfate reduction (ASR) and dissimilatory sulfate reduction (DSR). The dissimilatory sulfate reduction exists as a major reduction pathway in the species of the genus *Desulfovibrio* or *Pseudodesulfovibrio*, which involves the reduction of sulfate (SO_4_^2-^) to H_2_S (S^2-^). The reduction of sulfite (SO_3_^2-^) to H_2_S (S^2-^) mediated by dissimilatory sulfite reductase (DsrAB) is the most important part of the reduction process (7). As of now, more than 77 species of the genus *Desulfovibrio* or *Pseudodesulfovibrio* have been described (8, 9). *Desulfovibrio* or *Pseudodesulfovibrio* species have often been isolated ubiquitously in nature, mainly from freshwater, and marine anaerobic systems (10, 11). Many species of the genus *Desulfovibrio* have been isolated frequently from marine environments, including *Desulfovibrio senegalensis* (8), *Desulfovibrio frigidus* (12), *Desulfovibrio alkalitolerans* (13), *Desulfovibrio inopinatus* (14), and *Desulfovibrio bizertensis* (15). *Pseudodesulfovibrio* is a new genus originally proposed and reclassified of four species of the genus *Desulfovibrio* in 2016, and they all belong to the family *Desulfovibrionaceae* (16). Most species of the genus *Pseudodesulfovibrio* have been isolated from marine sediments, including *Pseudodesulfovibrio piezophilus* (10), *Pseudodesulfovibrio indicus* (16), *Pseudodesulfovibrio profundus* (17), *Pseudodesulfovibrio portus* (4), and *Pseudodesulfovibrio cashew* (9). The relatively high diversity and abundance of SRB have been reported in marine sediments, suggesting that SRB play a crucial role in the elemental cycles in marine sediments (18, 19). For example, the sulfur cycle in marine sediments is primarily driven by the dissimilatory sulfate reduction to sulfide by the anaerobic SRB (20).

Deep-sea cold seeps are widely distributed in the edges of continental shelves and mainly characterized by gas and liquid hydrocarbons from deep geological sources (21). The deep-sea cold seep is a very specific methane- and sulfate-rich environment, in which SRB account for approximately 5-25% of microbial biomass in the surface of sulfate-rich zones and up to approximately 30-35% in the sulfate-methane transition zone (19). The anaerobic oxidation of methane (AOM) is the primary process of complex cold seep ecosystems, which is conjointly managed by a consortium of anaerobic methane-oxidizing archaea (ANME) and SRB (22, 23). However, only a few deep-sea SRB strains have been reported, limiting our understanding of their characteristics (e.g., material metabolism, element cycling, and ecological role). Thus, the deep-sea cold seep is one of the best locations to study deep-sea sulfur cycle mediated by SRB, it is utmost precious to obtain the typical SRB to explore their real metabolisms happened in this special environment.

In this study, we report the high abundance of SRB in deep-sea cold seep sediments. We isolated an anaerobic representative of deep-sea SRB from the deep-sea subsurface sediment. Combining physiological and proteomic approaches, we confirmed that strain zrk46 had strong ability to metabolize sulfate, thiosulfate, and sulfite via the dissimilatory sulfate reduction, accompanied by auxiliary metabolic effects of heterodisulfide reductase, ferredoxin, and nitrate reduction associated proteins, in both laboratory and deep-sea *in situ* conditions. Lastly, the metatranscriptomic results also showed that deep-sea SRB performed the sulfate reduction in deep-sea environment, which was coupled to the methane oxidation of ANME.

## RESULTS AND DISCUSSION

### Cultivation and morphology of a novel deep-sea sulfate-reducing bacterium

To gain preliminary insights of SRB existing in the deep-sea cold seep, operational taxonomic units (OTUs) sequencing was firstly performed to detect the relative abundance in SSU rRNA gene tag sequencing of SRB present in the deep-sea cold seep sediments. The result showed that *Desulfobacteraceae*, *Desulfobulbaceae*, and *Desulfovibrionaceae* were the top three families in surface sediment (RPC, 0-10 cm), all of which were sulfate-reducing bacteria (Fig. 1A). The proportion of *Desulfovibrionaceae* accounted for 51.3% of the whole bacterial domain at the family level in sample ZC2 (90-110 cm) (Fig. 1B), suggesting SRB were dominant in deep-sea cold seep sediments, which is similar to other marine environments (19). The diversity and abundance of SRB have been relatively high in deep-sea sediments, implying the vital importance of SRB in deep-sea environment.

**Fig. 1.**
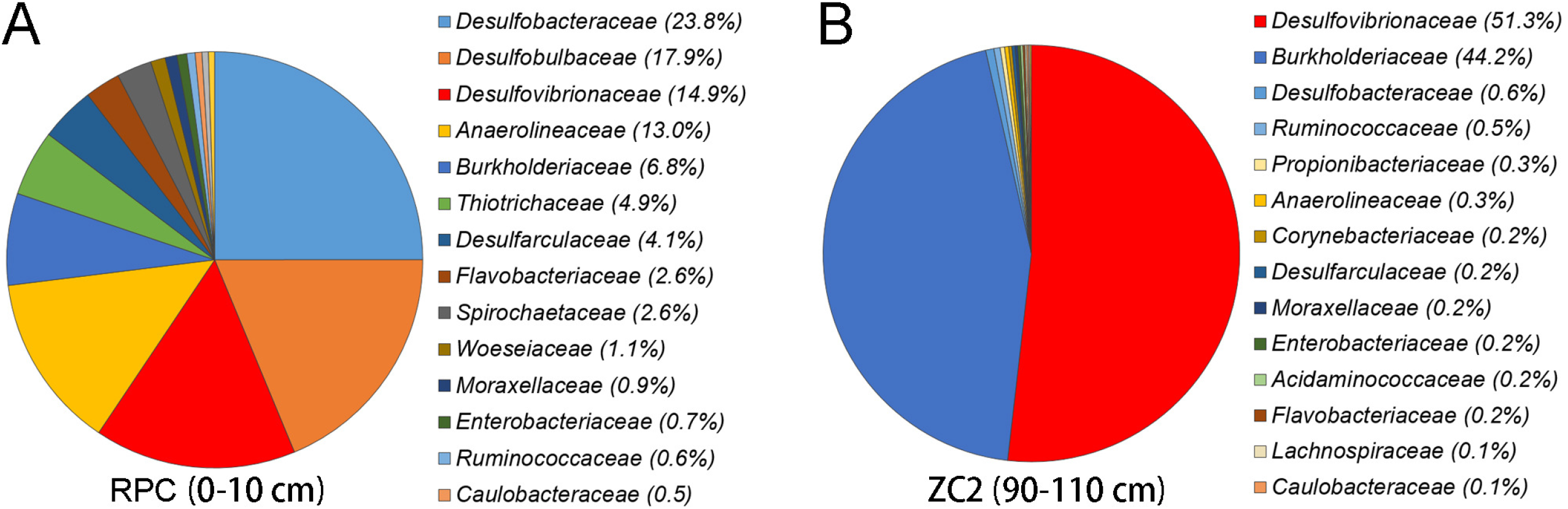
Detection of the abundance of SRB derived from deep-sea cold seep sediments. The community structure of two sampling sites (RPC and ZC2) in the cold seep sediments as revealed by 16S rRNA gene amplicon profiling. The relative abundances of OTUs representing different bacteria are shown at the family level.

To isolate SRB from deep-sea sediments, we developed an enrichment strategy by using a basal medium supplemented with 100 mM Na_2_SO_4_, which was often used as an electron acceptor for SRB. Then, these deep-sea sediment samples were anaerobically enriched at 28 °C for one month. Thereafter, the enriched cultures were plated on the solid medium in Hungate tubes, and individual colonies with distinct morphology were picked and cultured (Fig. 2A). Some of the cultured colonies were identified as SRB based on their 16S rRNA gene sequences. Among them, strain zrk46 possessed a fast growth rate and a complete purity, and then was chosen for further study. Under TEM observation, zrk46 showed a vibrioid or S-shaped, 2.0-4.5×0.3-0.7 µm in size, which was motile by means of a single polar flagellum (Fig. 2B).

**Fig. 2.**
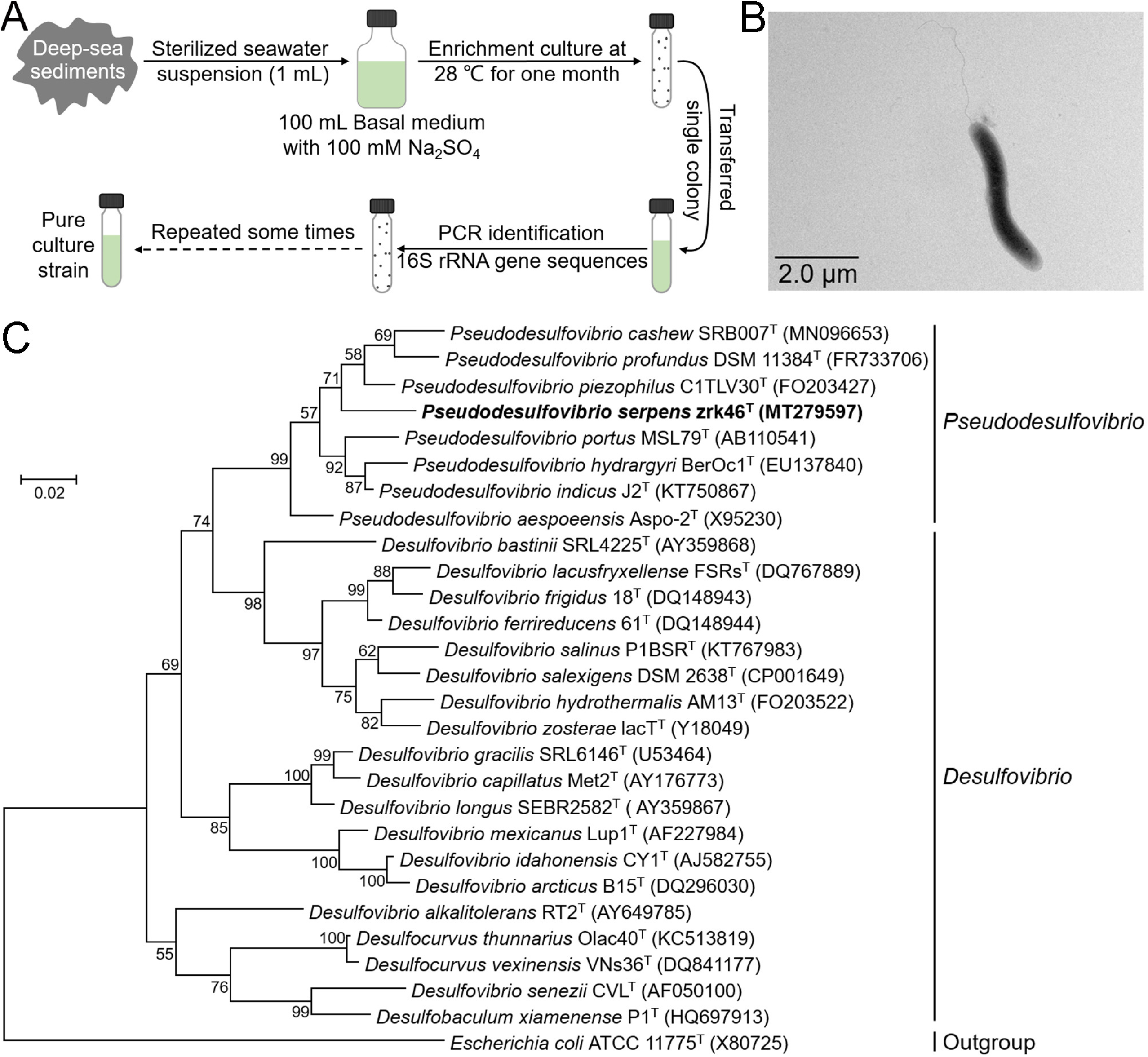
Isolation, morphology, and phylogenetic analyses of *Pseudodesulfovibrio serpens* zrk46. (A) Diagram showing the strategy used to isolate deep-sea SRB. (B) TEM observation of the morphology of strain zrk46. (C) Phylogenetic analysis of strain zrk46. Phylogenetic placement of strain zrk46 within the genus *Pseudodesulfovibrio*, based on almost complete 16S rRNA gene sequences. The NCBI accession number for each 16S rRNA gene is indicated after each corresponding strain’s name. The tree was inferred and reconstructed using the maximum likelihood criterion. Bootstrap values are based on 1000 replicates. The 16S rRNA gene sequence of *Escherichia coli* ATCC 11775^T^ was used as the outgroup. Bar, 0.02 substitutions per nucleotide position.

### Physiological characteristics, genome and phylogeny of strain zrk46

The detailed physiological characteristics of strain zrk46 and the type strain *Pseudodesulfovibrio profundus* DSM 11384^T^ were listed in Table S1. Strain zrk46 was grew at 0-80 g/L NaCl (optimum: 40 g/L) and grew at 16-45 °C (optimum 28 °C). The growth was observed at pH 6.0-8.5, the optimum being around 7.0. Compared with the type strain DSM 11384^T^, strain zrk46 showed its higher capacity to utilize different substrates as electron donors (fumarate, butyrate, formate, lactate, succinate, and malate) and acceptors (sulfate, sulfite, thiosulfate, and nitrate). The major fatty acids of strain zrk46 (>5 % of the total) were iso-C_15:0_ (28.53%), anteiso-C_15:0_ (9.66%), C_16:1_*cis*9 (6.17%), C_16:0_ (17.29%), iso-C_17:0_ (6.21%), C_18:1_*cis*11/*trans*9/*trans*6 (6.88%), and C_18:0_ (7.42%) (Table S2). The amount of iso-C_15:0_, C_16:1_*cis*9, C_16:0_, iso-C_17:0_, C_18:1_*cis*11/*trans*9/*trans*6, and C_18:0_ in strain zrk46 were higher than that found in strain DSM 11384^T^. However, the amount of anteiso-C_15:0_ was lower in strain zrk46 (9.66 %) than that found in strain DSM11384 (15.22 %). The major polar lipids in strain zrk46 were phosphatidylethanolamine (PE), diphosphatidylglycerol (DPG), phosphatidylglycerol (PG), aminophospholipid (APL), and two phospholipids (PL) (Fig. S1).

To assess genomic features of strain zrk46, its whole genome was sequenced and analyzed (Fig. S2). The chromosomal DNA G+C content of strain zrk46 was 53.26 %, and its genome size was 4,082,342 bp. Strain zrk46 does not possess any plasmid. Annotation of the genome of strain zrk46 revealed it consisted of 3717 predicted genes including 101 RNA genes (12 rRNA genes, 76 tRNA genes, and 13 other ncRNAs). To further clarify the phylogenetic position of strain zrk46, the genome relatedness values were calculated by the average nucleotide identity (ANI), *in silico* DNA-DNA similarity (*is*DDH), and the tetranucleotide signatures (Tetra), against the genomes of the four type strains of the genus *Pseudodesulfovibrio* (Table S3). The average nucleotide identities (ANIb) of strain zrk46 with strains Aspo-2^T^, J2^T^, C1TLV30^T^, and DSM 11384^T^ were 72.36%, 72.83%, 71.47%, and 73.31%, respectively. The average nucleotide identities (ANIm) of strain zrk46 with strains Aspo-2^T^, J2^T^, C1TLV30^T^, and DSM 11384^T^ were 83.43%, 83.63%, 83.85%, and 83.87%, respectively. The amino acid identities (AAI) of strain zrk46 with strains Aspo-2^T^, J2^T^, C1TLV30^T^, and DSM 11384^T^ were 74.5%, 73.9%, 76.4%, and 77.4%, respectively. The high AAI values to *Pseudodesulfovibrio* species identify strain zrk46 as a novel member of the genus *Pseudodesulfovibrio*. Based on digital DNA-DNA hybridization employing the Genome-to-Genome Distance Calculator GGDC (24), the *in silico* DDH estimates for zrk46 with strains Aspo-2^T^, J2^T^, C1TLV30^T^, and DSM 11384^T^ were 19.00%, 19.60%, 18.70%, and 19.90%, respectively. The tetranucleotide signatures (Tetra) of strain zrk46 with strains Aspo-2^T^, J2^T^, C1TLV30^T^, and DSM 11384^T^ were 0.75847, 0.73201, 0.75404, and 0.93565, respectively. These results demonstrated the genome of strain zrk46 to be clearly below established ‘cut-off’ values (ANIb: 95%, ANIm: 95%, AAI: 95%, *is*DDH: 70%, TETRA: 0.99) for defining bacterial species (25), suggesting strain zrk46 represented a novel species within the genus *Pseudodesulfovibrio* as currently defined.

To further confirm the taxonomy of strain zrk46, we performed phylogenetic analyses. The maximum likelihood tree of 16S rRNA indicated that strain zrk46 belonged to the genus *Pseudodesulfovibrio* (Fig. 2C). Based on the comparative 16S rRNA gene sequence analysis using BLAST tool and NCBI GenBank database, the 16S rRNA sequence similarity calculation using the NCBI server indicated that the closest relatives of strain zrk46 were *Pseudodesulfovibrio piezophilus* C1TLV30^T^ (96.11%), *Pseudodesulfovibrio aespoeensis* Aspo-2^T^ (95.27%), *Pseudodesulfovibrio profundus* DSM 11384^T^ (95.16%), and *Pseudodesulfovibrio indicus* J2^T^ (95.11%). Recently, the taxonomic threshold for species based on 16S rRNA gene sequence identity value was 98.65% (26). Together, based on phylogenetic, genomic, and phenotypic characteristics, we proposed that strain zrk46 might be a novel representative of the genus *Pseudodesulfovibrio*, for which the name *Pseudodesulfovibrio serpens* sp. nov. is proposed.

### Description of Pseudodesulfovibrio serpens sp. nov

*Pseudodesulfovibrio serpens* (ser’pens. L. fem. n. serpens [italic type] the snake).

Cells are Gram-stain-negative, strictly anaerobic, S-shaped or vibrioid, 2.0-4.5×0.3-0.7 µm in size, motile by a single polar flagellum. Catalase-positive, oxidase-positive. Fumarate, butyrate, formate, lactate, succinate, and malate are oxidized with sulfate reduction. Sulfate, sulfite, thiosulfate, and nitrate serve as electron acceptors. Growth is observed at salinities from 0 to 80 g/L NaCl (optimum: 40 g/L), from pH 6.0 to 8.5 (optimum 7.0) and at temperatures between 16 and 45 °C (optimum 28 °C). The major polar lipids in strain zrk46 are phosphatidylethanolamine (PE), diphosphatidylglycerol (DPG), phosphatidylglycerol (PG), aminophospholipid (APL), and two phospholipids (PL). Major fatty acids (>10 %) are iso-C_15:0_ (28.53 %) and C_16:0_ (17.29 %). The genome size of the type strain zrk46 is around 4.08 Mbp and the genomic DNA G+C content is 53.26 %.

The type strain, zrk46 (=MCCC 1K04422^T^), is isolated from the cold seep in the South China Sea. The GenBank 16S rRNA gene sequence accession number for isolate zrk46 is MT279597 and the corresponding GenBank NCBI accession number of the genome sequence (CP051216).

### Effects of Na_2_SO_4_, Na_2_S_2_O_3_, Na_2_SO_3_, and Na_2_S on *P. serpens* zrk46 growth

As SRB are essential anaerobic microorganisms mediating sulfur cycling (27), we firstly analyzed the sulfate reduction associated genes in the strain zrk46 genome. As expected, the genes encoding dissimilatory sulfate reduction closely related proteins were present in the genome of strain zrk46 (Fig. 3A). Inside the cytoplasm, sulfate is activated by the sulfate adenylyltransferase (Sat) to adenosine 5’-phosphosulfate (APS). Subsequently, APS is reduced to sulfite by the adenylylsulfate reductase (AprAB). Sulfite is a key intermediate in the dissimilatory sulfate reduction, which is reduced by the dissimilatory sulfite reductase (DsrAB) and a sulfur transfer protein (DsrC) together with the Dsr complex (DsrMKJOP) required for sulfate reduction (Fig. 3B) (28, 29). To date, most studies on SRB have been related to lake (30), oil reservoirs (31), freshwater sediments (32), soil (33), gut (34) and so on, with only few reports on the deep-sea extreme environment. Here, we tested the effects of different sulfur-containing substances (including Na_2_SO_4_, Na_2_S_2_O_3_, Na_2_SO_3_, and Na_2_S) on strain zrk46 growth. These assays showed that adding Na_2_SO_4_, Na_2_S_2_O_3_, or Na_2_SO_3_ to the culture medium increased strain zrk46 growth, while adding Na_2_S inhibited growth (Fig. 3C). Subsequently, we performed proteome sequencing analysis and found that most dissimilatory sulfate reduction-associated proteins were simultaneously upregulated in the presence of Na_2_SO_4_, Na_2_S_2_O_3_, and Na_2_SO_3_ (Fig. 3D). These results showed that strain zrk46 could effectively reduce sulfate, thiosulfate and sulfite to hydrogen sulfide via dissimilatory sulfate reduction that coupled to the oxidation of organic matter (35, 36), from which it can obtain energy to promote its growth.

**Fig. 3.**
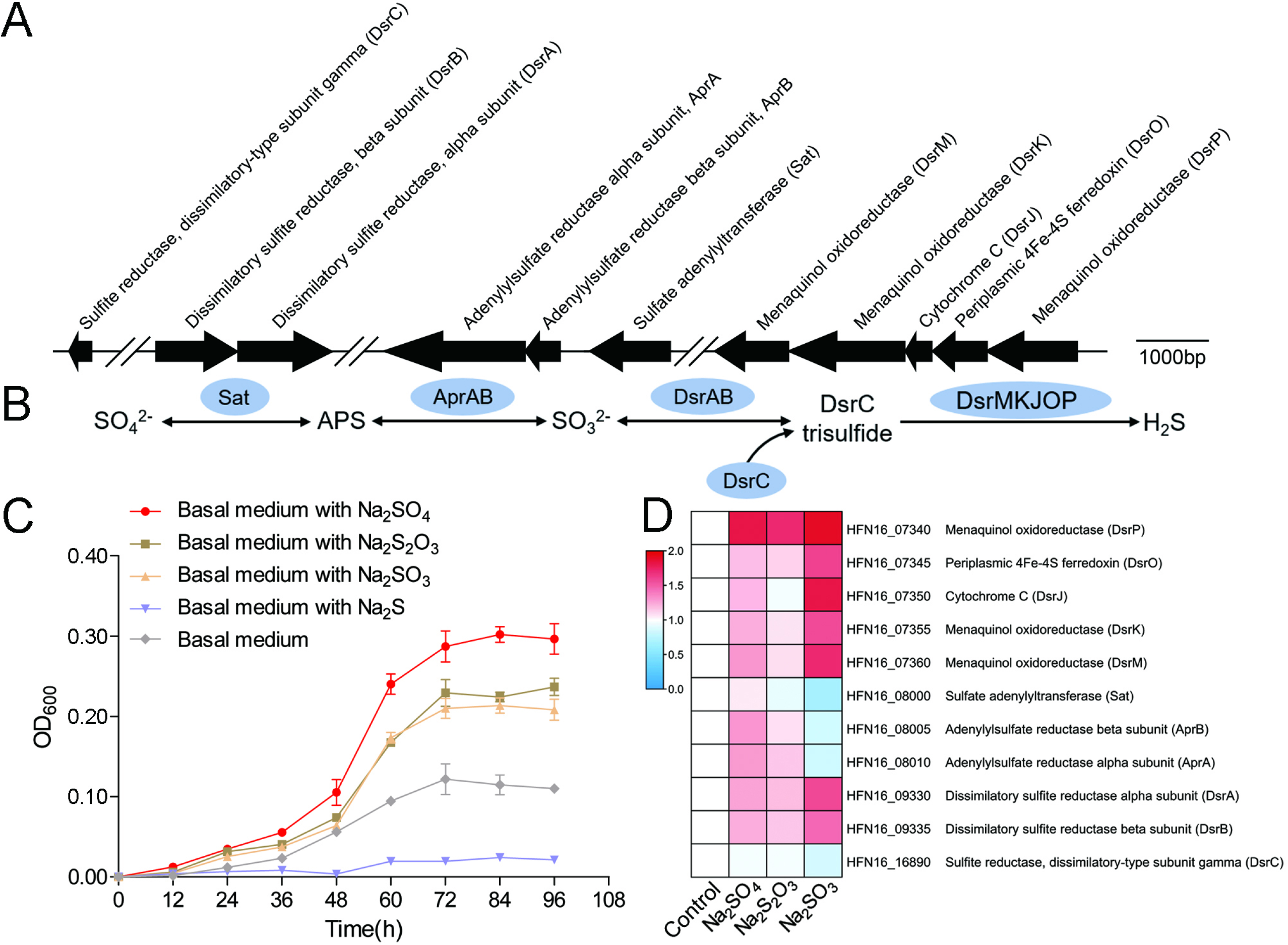
Sulfur metabolism assays of *Pseudodesulfovibrio serpens* zrk46. (A) The gene cluster containing the typical dissimilatory sulfate reductase operon and associated genes identified in the genome of strain zrk46. (B) Proposed pathway of dissimilatory sulfate reduction existing in strain zrk46. Sat, sulfuradenylyltransferase; APS, adenylyl sulfate; AprAB, adenylylsulfate reductase; DsrABC, reverse-type dissimilatory sulfite reductase; DsrMKJOP, sulfite reduction-associated complex. (C) Growth curves of strain zrk46 cultivated in basal medium alone and supplemented with either 20 mM Na_2_SO_4_, 20 mM Na_2_S_2_O_3_, 20 mM Na_2_SO_3_, or 5 mM Na_2_S. (D) Proteomics-based heat map showing the relative expression levels of proteins associated with sulfur metabolism.

Moreover, we observed that the expression of heterodisulfide reductase related proteins were simultaneously upregulated in the presence of Na_2_SO_4_, Na_2_S_2_O_3_, and Na_2_SO_3_ (Fig. 4A, B, and C). Heterodisulfide reductase was widespread in the domains Bacteria and Archaea, and play roles in a greater diversity of energy-conserving metabolisms, including the reduction of sulfate (37, 38) and the oxidation of inorganic sulfur compounds (39). Therefore, the result indicated that heterodisulfide reductase could assist the sulfate reduction process of strain zrk46. Interestingly, the expression of nitrogen metabolism associated proteins (including nitrate reductase, nitrite reductase, nitroreductase) were simultaneously upregulated in the presence of Na_2_SO_4_, Na_2_S_2_O_3_, and Na_2_SO_3_ (Fig. 4A, B, and C), which indicate that these reductases might play a role in the sulfate reduction process of strain zrk46, which need further studies. Although they cannot directly affect inorganic sulfur compounds, they might contribute significantly to the electron transfer and formation energy of strain zrk46. Alternatively, the expression of iron–sulfur (Fe-S) proteins and ferredoxins were also upregulated in the presence of Na_2_SO_4_, Na_2_S_2_O_3_ and Na_2_SO_3_ (Fig. 4D, E, and F). Ferredoxins comprise a large family of Fe–S proteins that transfer electrons in diverse biological processes (40). The type Fe-S proteins in strain zrk46 was 2Fe-2S cluster and 4Fe-4S cluster, but only the expressions of 4Fe-4S associated proteins were upregulated, indicating that it played a functional role in electron transfer processes relevant for the sulfate reduction process of strain zrk46. In the future, the genetic operating system of this strict anaerobic bacterium needs to be constructed to further confirm its function.

**Fig. 4.**
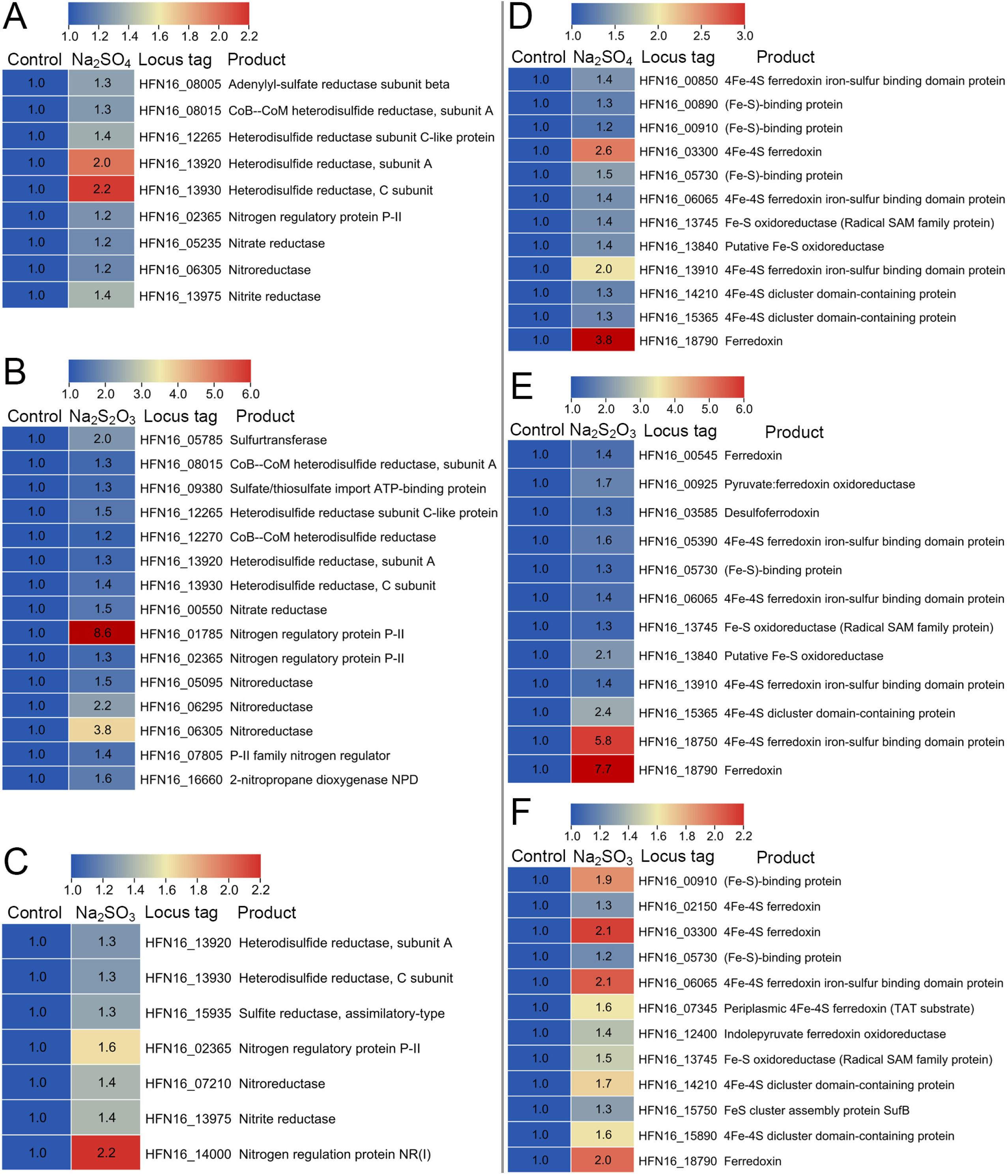
Proteomic analysis of *Pseudodesulfovibrio serpens* zrk46 cultivated in basal medium alone and supplemented with either 20 mM Na_2_SO_4_, 20 mM Na_2_S_2_O_3_, or 20 mM Na_2_SO_3_. Proteomics-based heat map showing the upregulated proteins associated with sulfur metabolism and nitrogen metabolism (A, B, and C). Proteomics-based heat map showing the upregulated proteins associated Fe-S proteins (D, E, and F). “Control” indicates basal medium. “Na_2_SO_4_, Na_2_S_2_O_3_ and Na_2_SO_3_” indicate basal medium supplemented with 20 mM Na_2_SO_4_, 20 mM Na_2_S_2_O_3_, and 20 mM Na_2_SO_3_, respectively.

### Proteomic analysis of *P. serpens* zrk46 cultured in deep-sea conditions

Considering stain zrk46 was isolated from the deep-sea environment, we next sought to explore its actual metabolisms performed in the deep-sea environment. We thus performed the *in situ* cultivation of strain zrk46 in anaerobic bags either without or with exposure to the deep-sea environment (where we isolated this bacterium) for ten days, as previously described (41). Subsequently, the strain zrk46 cells were collected and performed the proteomic analyses. The upregulated proteins were predominantly mapped to COG categories representing energy production and conversion, amino acid transport and metabolism, cell motility, and signal transduction mechanisms (Fig. 5A). Interestingly, the proteomic results clearly showed that the expressions of thiosulfate reductase, heterodisulfide reductase, and sulfite reductase were upregulated when compared to that cultured under the closed condition (Fig. 5B), which were consistent with the laboratory conditions, indicating dissimilatory sulfate reduction indeed happened in the deep sea. The expressions of nitrate reductase and nitroreductase were also upregulated, which indicated that strain zrk46 performed active nitrogen metabolism in the deep sea, or it could assist strain zrk46 to acquire electrons from electron transfer chains to reduce sulfate, which awaits future studies (42). Consistently, the expressions of most 4Fe-4S proteins and ferredoxins were upregulated in the deep sea (Fig. 5C), indicating that they indeed play a vital role in the deep-sea environment, possibly involved in electron transfer during the sulfate reduction of strain zrk46. In addition, the formate dehydrogenases were also highly expressed in the deep-sea environment (Fig. 5D), suggesting that strain zrk46 could enhance formate metabolism and thus providing more electrons and protons (reducing power) for the sulfate reduction process (43). In anoxic environments, SRB are primarily responsible for organic carbon oxidation, because sulfate is often the predominant electron acceptor (27, 44). It has previously been reported that sulfate reduction could facilitate the organic matter oxidation up to 50% in the marine sediments (45). Notably, most ABC transporters (associated with amino acids, sugars, and ions etc.) were upregulated (Fig. 5E), which indicating that strain zrk46 could effectively ingest and degrade the organic compounds coupled sulfate reduction process in the deep-sea environment (35, 36). In conclusion, we performed the *in situ* experiments on deep-sea SRB to reveal its true metabolic characteristics in deep-sea environment.

**Fig. 5.**
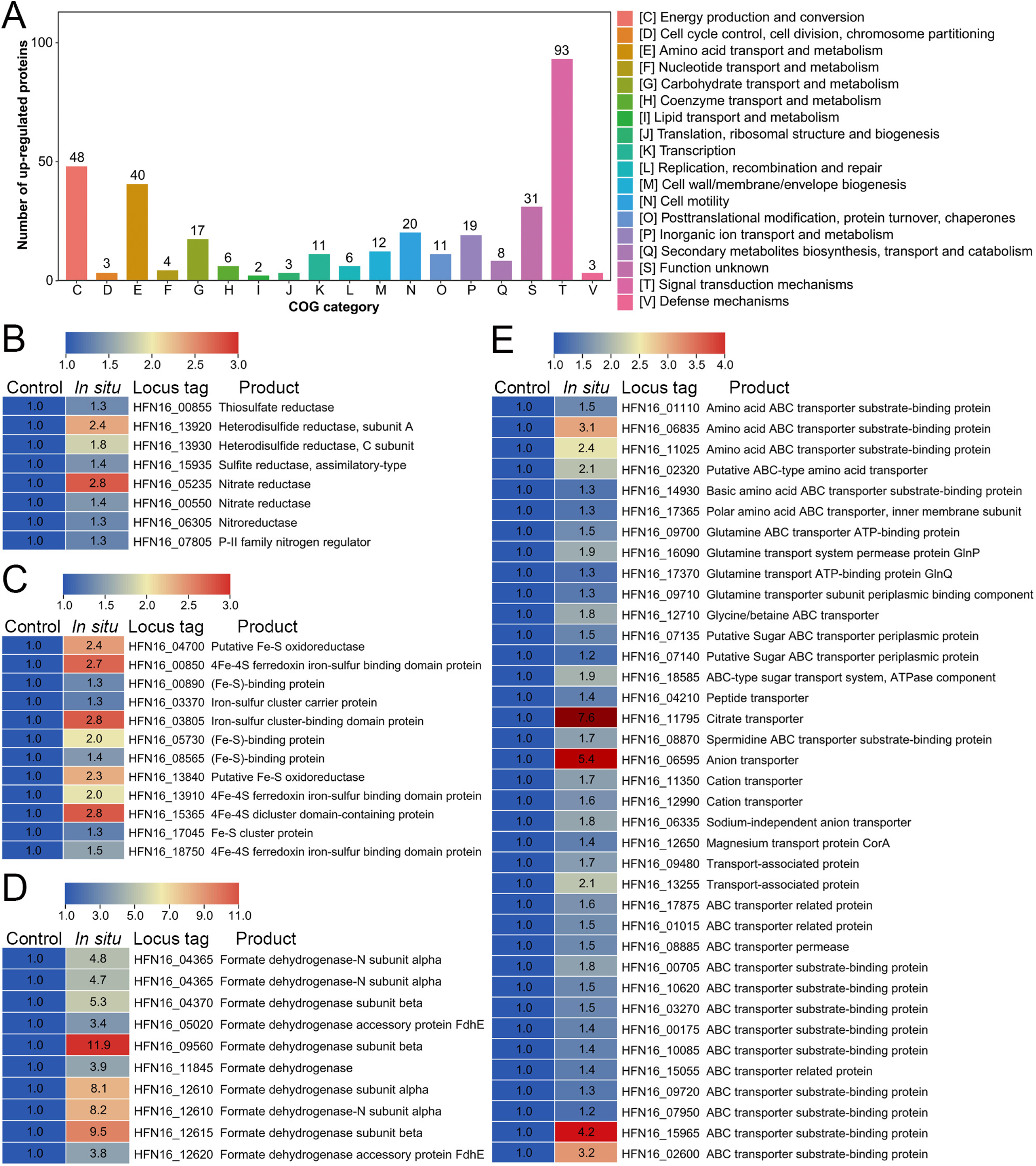
Proteomic analysis of *Pseudodesulfovibrio serpens* zrk46 incubated in the deep-sea cold seep. (A) COG category showing the significant enrichment of upregulated proteins in strain zrk46 cultivated in the deep-sea cold seep. Proteomics-based heat map showing the upregulated proteins associated with sulfur metabolism and nitrogen metabolism (B), Fe-S proteins (C), formate dehydrogenase (D), and ABC transporter proteins (E). “Control” indicates strain zrk46 cultured in deep sea condition without exchanging with outside; “*In situ*” strain zrk46 cultured in deep sea condition exchanging with outside.

### *In situ* metatranscriptomic analysis of metabolism of deep-sea SRB and its association with anaerobic methane-oxidizing archaea (ANME)

Given the high abundance of SRB and the real sulfur metabolism of strain zrk46 in the deep-sea cold seep, we performed the metatranscriptomic sequencing analysis to investigate real metabolisms of other deep-sea SRB. The result showed that the genes encoding adenosine-5’-phosphosulfate (APS) reductase, adenylylsulfate reductase, sulfate adenylyltransferase, thiosulfate reductase, dissimilatory sulfite reductase, and sulfite exporter of different SRB were upregulated in the cold seep sediment compared to the sediment that far away from the cold seep (Fig. 6A). This result indicates that SRB perform the sulfate reduction in deep-sea environment and play a pivotal role in the sulfur biogeochemical cycling (46). In contrast, the genes encoding nitrate reductase and nitrite reductase of most SRB were downregulated, while only a few were up-regulated (Fig. 6B), possibly due to the high concentration of sulfate in the deep-sea cold seep. Given that a consortium of anaerobic methane-oxidizing archaea (ANME) and SRB can conjointly operate the anaerobic oxidation of methane (AOM) (23, 47), we analyzed the expression of genes associated with methane metabolism in ANME via the metatranscriptome. The result showed that the genes encoding methyl coenzyme M reductase, CoB-CoM heterodisulfide reductase, formate dehydrogenase, H4MPT-linked C1 transfer pathway protein, and F_420_H_2_ dehydrogenase of ANME-2 cluster archaeon were all upregulated, which were involved in the oxidation of methane (Fig. 7A). In addition, the genes encoding *c*-type cytochromes in SRB and ANME-2 cluster archaeon were all upregulated (Fig. 7B). Recent evidence suggests that electrons are transferred from ANME archaea to SRB partners using outer membrane c-type cytochromes (48, 49), which might conduct direct interspecies electron transfer between SRB and ANME-2 in the deep-sea environment (50, 51). Moreover, we found that the concentrations of methane and sulfate in this cold seep sediment were 2642 μM and 28 mM (Table S4), respectively, which were sufficient to support the processes of sulfate reduction and methane oxidation. Therefore, we speculate that deep-sea SRB may be couple to the methane oxidation of ANME-2 in the process of sulfate reduction, while whether nitrate reduction in SRB is coupled to methane oxidation of ANME-2 remains to be confirmed. According to the close relationship between ANME and SRB, we are co-culturing strain zrk46 together with the enrichment of ANME and hoping to increase the abundance of ANME and even obtain its pure culture in the future.

**Fig. 6.**
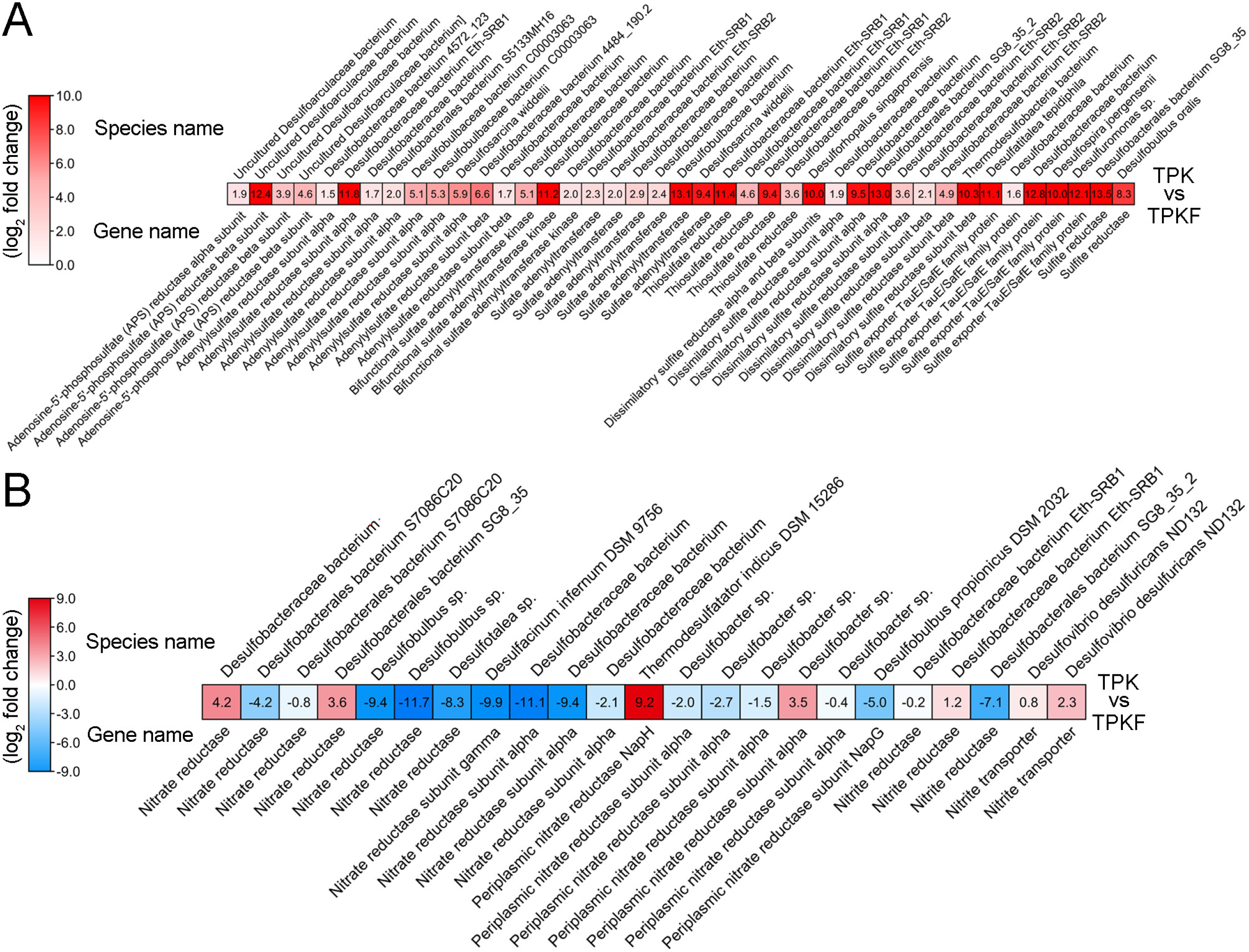
*In situ* metatranscriptomic analysis of the metabolic characteristics of deep-sea SRB. The relative expression levels of genes encoding proteins associated with sulfur metabolism (A) and nitrogen metabolism (B) in deep-sea SRB. The numbers in panels A and B represent the fold change of gene expression (by using the log_2_ value). TPK, sedimentl sample from the cold seep vent; TPKF, sedimental sample far away from the cold seep vent.

**Fig. 7.**
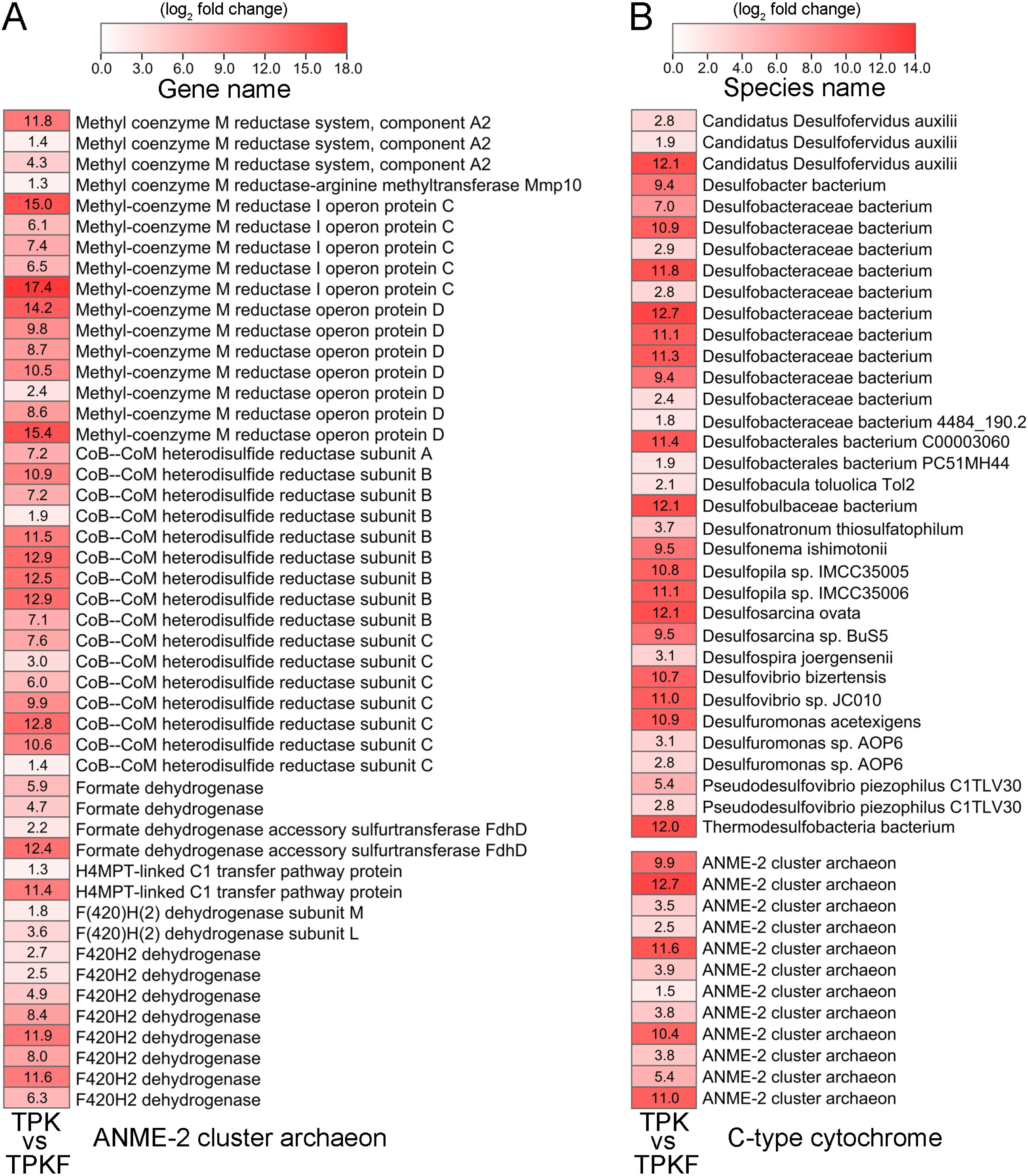
*In situ* metatranscriptomic analysis of the expression of anaerobic methane oxidation related genes of ANME-2 cluster archaeon (A) and the genes encoding c-type cytochromes of SRB and ANME (B). The numbers represent the fold change of gene expression (by using the log_2_ value). TPK, sedimentl sample from the cold seep vent; TPKF, sedimental sample far away from the cold seep vent.

## CONCLUSION

The integrative analysis of genomic, physiological, laboratory and *in situ* proteomic, and metatranscriptomic data in the present study identified the sulfur metabolic characteristics of a novel deep-sea sulfate-reducing bacterium (zrk46) in both laboratory and deep-sea *in situ* environments. These findings expand our understanding of deep-sea SRB, while highlighting their importance for driving deep-sea elemental cycles. Important information is also provided regarding the sulfate reduction of deep-sea SRB in deep-sea environment might be coupled to the methane oxidation of ANME-2 through direct interspecies electron transfer via cytochromes.

## MATERIALS AND METHODS

### Sampling and OTUs analysis

The deep-sea sediment samples were collected by *RV KEXUE* from a typical cold seep in the South China Sea (E 119°17’07.322’’, N 22°06’58.598’’) at a depth of approximately 1,146 m in July of 2018. We selected two sedimentary samples (RPC and ZC2 at depth intervals of 0-10 and 90-110 cm, respectively) for OTUs sequencing that performed by Novogene (Tianjin, China). Briefly, total DNAs from these samples were extracted by a CTAB/SDS method (52) and diluted to 1 ng/µL with sterile water, and then used for PCR template. We used the primers (341F: 5’-CCTAYGGGRBGCASCAG and 806R: 5’-GGACTACNNGGGTATCTAAT) to amplify the 16S rRNA genes of distinct regions (V3/V4). Then, we used a Qiagen Gel Extraction Kit (Qiagen, Germany) to purify these PCR products for libraries construction. Sequencing libraries were generated using TruSeq® DNA PCR-Free Sample Preparation Kit (Illumina, USA) following the manufacturer’s instructions, and then assessed on the Qubit@ 2.0 Fluorometer (Thermo Scientific, USA) and Agilent Bioanalyzer 2100 system. The library was sequenced on an Illumina NovaSeq platform and 250 bp paired-end reads were generated and merged using FLASH (V1.2.7, http://ccb.jhu.edu/software/FLASH/) (53). Quality filtering on raw tags was performed according to the QIIME (V1.9.1, http://qiime.org/scripts/split_libraries_fastq.html) quality controlled process to obtain the high-quality clean tags (54). To detect chimera sequences (55), the tags were compared with the reference database (Silva database version 138, https://www.arb-silva.de/) using UCHIME algorithm (UCHIME Algorithm, http://www.drive5.com/usearch/manual/uchime_algo.html) (56). Sequence analyses were performed by Uparse software (Uparse v7.0.1001, http://drive5.com/uparse/) (57) and the sequences with ≥97% similarity were assigned to the same OTUs. The representative sequence for each OTU was screened for further annotation. Finally, we used the Silva Database (http://www.arb-silva.de/) (58) with the Mothur algorithm to annotate taxonomic information.

### Enrichment and cultivation of deep-sea SRB

To isolate and cultivate deep-sea SRB, 1.0 g deep-sea sediment sample of RPC or ZC2 was suspended with 1 mL sterilized seawater, respectively, and then added to a 250 mL anaerobic bottle containing 100 mL inorganic medium (containing 2.0 g/L sodium lactate, 1.0 g/L NH_4_Cl, 1.0 g/L NaHCO_3_, 1.0 g/L CH_3_COONa, 0.5 g/L KH_2_PO_4_, 0.2 g/L MgSO_4_.7H_2_O, 0.7 g/L cysteine hydrochloride, 500 µL/L 0.1 % (w/v) resazurin, pH 7.0) supplemented with 100 mM Na_2_SO_4_ under a 100% N_2_ atmosphere. The medium was prepared under a 100% N_2_ gas phase and sterilized by autoclaving at 115 °C for 30 minutes. The inoculated media were anaerobically incubated at either 4 °C or 28 °C for one month. The inorganic medium supplemented with 100 mM Na_2_SO_4_ and 15 g/L agar was evenly spread on to the inside wall of a Hungate tube, which formed a thin layer of medium for the bacteria to grow. After this, 100 µL of the enriched culture was anaerobically transferred into the anaerobic roll tubes and then spread on to the medium layer. These tubes were also anaerobically cultured for three days. Individual colonies growing at 28 °C were selected using sterilized bamboo sticks according to distinct morphology (such as size, color, etc); they were then cultured in the 15 mL Hungate tube containing 10 mL inorganic medium supplemented with 100 mM Na_2_SO_4_ at 28 °C for three days under a 100% N_2_ atmosphere. Thereafter, the amplification and sequencing of 16S rRNA genes were performed to identify these cultures. The primers 27F (5′-AGAGTTTGATCCTGGCTCAG-3′) and 1492R (5′-GGTTACCTTGTTACGACTT-3′) were used to amplify 16S rRNA gene sequences. PCR conditions were set as following: pre-denaturation at 95 °C for 10 minutes; denaturation at 95 °C for 15 seconds, annealing at 54 °C for 15 seconds, extension at 72 °C for 20 seconds, in 30 cycles; and final extension at 72 °C for 5 minutes. These PCR amplification products were sequenced in Tsingke Biotechnology Co., Ltd (Beijing, China), and the sequences were analyzed by BLASTn of the NCBI database. One strain (zrk46) was identified as a member of the genus *Pseudodesulfovibrio*, but was noted to have less than 97% 16S rRNA gene sequence similarity to other cultured strains; strain zrk46 was therefore selected and purified by repeating the Hungate roll-tube method. The purity of strain zrk46 was confirmed regularly by repeating partial sequencing of the 16S rRNA gene and by observation using a transmission electron microscope (TEM). Since strain zrk46 grew slowly in the inorganic medium, we added extra organic substances in the medium as following: 1.0 g/L yeast extract, 1.0 g/L peptone, 2.0 g/L sodium lactate, 1.0 g/L NH_4_Cl, 1.0 g/L NaHCO_3_, 1.0 g/L CH_3_COONa, 0.5 g/L KH_2_PO_4_, 0.2 g/L MgSO_4_.7H_2_O, 0.7 g/L cysteine hydrochloride, 500 µL/L 0.1 % (w/v) resazurin, pH 7.0, and the corresponding medium was named ‘basal medium’ in this study.

### TEM observation

To observe the morphological characteristics of strain zrk46, 10 mL culture was collected by centrifuging at 4000 × *g* for 5 minutes. Cells were then washed three times with PBS buffer (137 mM NaCl, 2.7 mM KCl, 10 mM Na_2_HPO_4_, 1.8 mM KH_2_PO_4_, 1 L sterile water, pH 7.4). Finally, the cells were suspended in 20 μL PBS buffer, and then transferred onto copper grids coated with a carbon film by immersing the grids in the cell suspension for 30 minutes (41). All samples were examined under TEM (HT7700, Hitachi, Japan).

### Genome sequencing and analysis

Genomic DNA of strain zrk46 was extracted from 0.5 L cells that cultured for three days at 28 °C. The DNA library was prepared using the Ligation Sequencing Kit (SQK-LSK109, UK), and sequenced using a FLO-MIN106 vR9.4 flow-cell for 48 hours on MinKNOWN software v1.4.2 (Oxford Nanopore Technologies, UK). The detailed sequencing procedures were performed as previously described (41). Moreover, the genome relatedness values were calculated by several approaches: amino acid identity (AAI), average nucleotide identity (ANI) based on the MUMMER ultra-rapid aligning tool (ANIm), ANI based on the BLASTN algorithm (ANIb), the tetranucleotide signatures (TETRA), and *in silico* DNA–DNA similarity (*is*DDH). The amino acid identity (AAI) values were calculated by AAI-profiler (http://ekhidna2.biocenter.helsinki.fi/AAI/) (59). ANIm, ANIb, and TETRA frequencies were calculated by the JSpecies WS (http://jspecies.ribohost.com/jspeciesws/) (60). The *is*DDH values were calculated by the Genome-to-Genome Distance Calculator (GGDC) (http://ggdc.dsmz.de/) (24). The *is*DDH results were based on the recommended formula 2, which is independent of genome size. The recommended species criterion cut-offs were used: 95% for the AAI, ANIb and ANIm, 0.99 for the TETRA signature, and 70% for the *is*DDH (25).

### Phenotypic characteristics analyses

Unless stated otherwise, physiological characterization was carried out anaerobically in basal medium supplemented with 20 mM Na_2_SO_4_. Growth was tested at different temperatures (4, 16, 28, 30, 32, 37, 45, 60, 70, 80 °C) for 10 days. The pH range for growth was tested from pH 4.0 to pH 10.0 with increments of 0.5 pH units. The pH of the culture medium was adjusted by 6 M HCl for low pH and 10 % NaHCO_3_ (w/v) for high pH. Salt tolerance was tested on the modified basal medium (replaced sea water with distilled water) supplemented with 0-10.0% (w/v) NaCl (1% intervals) for ten days. Determination of electron donors and acceptors for growth were performed with different electron donors at 20 mM (acetate, fumarate, formate, pyruvate, lactate, malate, methanol, fructose, propionate, butyrate, succinate, glycine, ethanol) and different electron acceptors: elemental sulfur (1%, w/v), sulfate (20 mM), sulfite (20 mM), thiosulfate (20 mM), nitrate (20 mM), and nitrite (20 mM). For each substrate, three biological replicates were performed.

For chemotaxonomic analysis, cells of zrk46 were cultured and collected under the same conditions unless stated otherwise, with the closely related type strain (*Pseudodesulfovibrio profundus* DSM 11384^T^) were grown on basal solid medium for 4 d at 28 °C under the same condition. Cells of zrk46 and the closely related type strain were harvested from cultures at the mid-exponential phase of growth and freeze-dried. Cellular fatty acids were extracted and determined from dried cells by using GC (model 7890A, Agilent, USA) according to the protocol of the Sherlock Microbial Identification System (61). Polar lipids were extracted and determined as described by Tindall *et al* (62).

### Phylogenetic analysis

To construct a maximum likelihood 16S rRNA phylogenetic tree, the full-length 16S rRNA gene sequences of strain zrk46 and other related taxa were obtained from the NCBI database (www.ncbi.nlm.nih.gov/). The phylogenetic tree was constructed using the W-IQ-TREE web server (http://iqtree.cibiv.univie.ac.at) (63) with the “GTR+F+I+G4” model, and the Interactive Tree of Life (iTOL v5) online tool (64) was used to edit the phylogenetic trees.

### Growth assays of strain zrk46

To assess the effects of different inorganic sulfur sources (20 mM Na_2_SO_4_, 20 mM Na_2_S_2_O_3_, 20 mM Na_2_SO_3_, and 5 mM Na_2_S) on strain zrk46 growth, we used a basal medium supplemented with the sulfur sources mentioned above. For each growth assay, 1 mL of strain zrk46 culture was inoculated in a 250 mL Hungate bottle containing 100 mL of the respective media. All Hungate bottles were anaerobically incubated at 28 °C. Bacterial growth was monitored by measuring OD_600_ values every 12 hours via a microplate reader until cell growth reached a stationary phase. Three replicates were performed for each condition.

### Proteomic analysis of sulfur metabolism of strain zrk46 cultured in laboratory condition

For proteomic analysis, cells suspension of strain zrk46 cultured in 200 mL of either basal medium or basal medium supplemented with different sulfur compounds (20 mM Na_2_SO_4_, 20 mM Na_2_S_2_O_3_, or 20 mM Na_2_SO_3_) at 28 °C for four days were collected at 8,000 ×*g* for 20 minutes. Subsequently, the cells were collected and sonicated three times on ice using a high intensity ultrasonic processor in lysis buffer (8 M urea, 1% Protease Inhibitor Cocktail). The remaining debris was removed by centrifugation at 12,000 ×*g*, 4 °C for ten minutes. Finally, the supernatant was collected and the protein concentration was determined with a BCA kit (Solarbio, China) according to the instructions. Proteomic sequencing analysis was performed by PTMBiolabs (Hangzhou, China), and the detailed protocols of proteomic sequencing technology were described in the Supplementary information.

### Proteomic analysis of sulfur metabolism of strain zrk46 cultured in the deep sea

To explore the actual metabolic characteristics of strain zrk46 conducted in the deep-sea cold seep, the *in situ* cultivation was performed. Briefly, 2 mL freshly incubated strain zrk46 cells was respectively transferred to three non-transparent anaerobic bags (which not allowing any exchanges between inside and outside; Aluminum-plastic composite film, Hede, China) with 200 mL basal medium each and set as control groups; on the other hand, 2 mL freshly incubated strain zrk46 cells was respectively transferred into three dialysis bags (8,000-14,000 Da cutoff, which allowing the exchanges of substances smaller than 8,000 Da but preventing bacterial cells from entering or leaving the bag; Solarbio, China) with 200 mL basal medium each and set as experimental groups. In June 2021, all the samples were placed simultaneously in the deep-sea cold seep (where strain zrk46 was isolated) for 10 days during the cruise of *Kexue* vessel. After 10 days *in situ* cultivation, cells of strain zrk46 were immediately collected and kept in the −80 °C freezer for future analysis. Before the proteomic sequencing analysis, the cells were checked by 16S rRNA gene sequencing to confirm the purity. The detailed protocol for proteomic sequencing was performed as described above.

### Metatranscriptomic analysis of metabolism of deep-sea SRB and its association with ANME

To explore the actual metabolic characteristics of SRB conducted in the deep-sea cold seep, the *in situ* metatranscriptomic analysis was performed. Two cold seep sediment samples (TPK, cold seep vent area; TPKF, far away from cold seep vent area) were selected for metatranscriptomic sequencing analysis in Shanghai Biozeron Biothchnology Co., Ltd. (Shanghai, China). Total RNAS were extracted from these sediments using TRIzol® Reagent according the manufacturer’s instructions and genomic DNA was removed using DNase I (TaKara, Japan). Then RNA quality was determined using 2100 Bioanalyser (Agilent, USA) and quantified using the ND-2000 (NanoDrop Technologies, USA). High-quality RNA sample (OD260/280=1.8∼2.2, OD260/230 ≥ 2.0, RIN ≥ 6.5, 28S:18S ≥ 1.0, >10μg) is used to construct sequencing library. The detailed protocols of library preparation, Illumina Hiseq sequencing, reads quality control and mapping, metatranscriptome assembly and annotation, and data analyses are described in the Supplementary information.

## Data availability

The full-length 16S rRNA gene sequence of strain zrk46 has been deposited at GenBank under the accession number MT279597. The complete genome sequence of strain zrk46 has been deposited at GenBank under the accession number CP051216. The mass spectrometry proteomics data have been deposited to the Proteome Xchange Consortium with the dataset identifier PXD043318. The raw amplicon sequencing data have been deposited to NCBI Short Read Archive (accession numbers: PRJNA675395). The raw metatranscriptomic sequencing data have been deposited to NCBI Short Read Archive (accession number: PRJNA988008).

## ACKNOWLEDGEMENTS

This work was funded by the Science and Technology Innovation Project of Laoshan Laboratory (Grant No. LSKJ202203103; 2022QNLM030004-3), the NSFC Innovative Group Grant (No. 42221005), Shandong Provincial Natural Science Foundation (ZR2021ZD28), Major Research Plan of the National Natural Science Foundation (Grant No. 92051107), Strategic Priority Research Program of the Chinese Academy of Sciences (Grant No. XDA22050301), Key deployment projects of Center of Ocean Mega-Science of the Chinese Academy of Sciences (Grant No. COMS2020Q04), and Key Collaborative Research Program of the Alliance of International Science Organizations (Grant No. ANSO-CR-KP-2022-08), and the Taishan Scholars Program (Grant No. tstp20230637) for Chaomin Sun. The authors are grateful to the captain and crew of the R/V *KEXUE*, as well as the *FAXIAN* ROV team for assistance with sample collection.

## AUTHOR CONTRIBUTIONS

CW and CS conceived and designed the study; CW conducted most of the experiments; RZ helped to isolate strain zrk46; CW and CS lead the writing of the manuscript; all authors contributed to and reviewed the manuscript.

